# iGenomics: Comprehensive DNA Sequence Analysis on your Smartphone

**DOI:** 10.1101/2020.02.11.944132

**Authors:** Aspyn Palatnick, Bin Zhou, Elodie Ghedin, Michael C. Schatz

**Affiliations:** Cold Spring Harbor High School, Cold Spring Harbor, NY 11724; Simons Center for Quantitative Biology, Cold Spring Harbor Laboratory, Cold Spring Harbor, NY 11724; University of Pennsylvania, Philadelphia, PA 19104; Department of Biology, New York University, New York, NY 10003; Department of Epidemiology, School of Global Public Health, New York, NY 10003; Departments of Computer Science and Biology, Johns Hopkins University, Baltimore MD, 21211

**Keywords:** speciation, bacteria, periodic selection, bacterial diversity

## Abstract

Following the miniaturization of integrated circuitry and other computer hardware over the past several decades, DNA sequencing is following a similar path. Leading this trend is the Oxford Nanopore sequencing platform, which currently offers the hand-held MinION instrument and even smaller instruments on the near horizon. This technology has been used in several important applications, including the analysis of genomes of major pathogens in remote stations around the world. However, despite the simplicity of the sequencer, an equally simple and portable analysis platform is not yet available.

iGenomics is the first comprehensive mobile genome analysis application, with capabilities to align reads, call variants, and visualize the results entirely on an iOS device. Implemented in Objective-C using the FM-index, banded dynamic programming, and other high-performance bioinformatics techniques, iGenomics is optimized to run in a mobile environment. We benchmark iGenomics using a variety of real and simulated Nanopore sequencing datasets and show that iGenomics has performance comparable to the popular BWA-MEM/Samtools/IGV suite, without needing a laptop or server cluster. iGenomics is available open-source (https://github.com/stuckinaboot/iGenomics) and for free on Apple’s App Store (https://apps.apple.com/us/app/igenomics-mobile-dna-analysis/id1495719841).

## Background

DNA sequencing technology has made tremendous progress over the past 30 years (Goodwin, McPherson, and McCombie 2016). The earliest automated approaches, beginning with the capillary-based Sanger sequencing devices in the 1980s, were large bench-top instruments requiring extensive sequencing facilities to prepare and sequence the DNA. In the 2000s, high throughput second-generation sequencing instruments advanced the field with more compact and simpler designs. However, these advances have been limited in their reach, because they are not readily accessible by most individual laboratories and citizen scientists.

Within the past few years, Oxford Nanopore Technologies (ONT, Oxford, UK) has introduced small inexpensive hand-held sequencing instruments that have made it possible to perform genomics experiments with minimal facilities and in essentially any environment. Nanopore sequencing technology works by measuring the change in ionic current as a DNA molecule is passed through a nanopore. The DNA molecules are typically a few hundred to tens of thousands of nucleotides long, sampled from random positions throughout the genome. Once sequenced, the raw signal data are base-called into nucleotide strings called reads (Wick, Judd, and Holt 2019), which are stored in fastq format and saved for further processing, especially read alignment and variant analysis.

Several algorithms are available for this analysis. Modern aligners, such as Bowtie (Langmead et al. 2009) or BWA-MEM (Li 2013), often use the Burrows-Wheeler Transform (BWT) (Burrows and Wheeler 1994) and the closely related FM-index (Ferragina and Manzini 2000) as their core indexing data structure. These new approaches are suited to large data sets because of their compact space requirements and fast alignment times. After alignment, variant calling platforms, such as Samtools (Li et al. 2009) or GATK (McKenna et al. 2010), systematically scan the alignments to find well supported variants in the sample using a statistical model to distinguish homozygous from heterozygous variants and rule out spurious sequencing errors. After this automated variant identification, priority variants are also often manually inspected using IGV (Robinson et al. 2011) and other genome browsers to review the evidence for the variant calls and further rule out false positives.

The standard approach for analyzing reads is to align the reads to a reference genome on high-end laptops, servers, or even supercomputers. While this is possible for those with access to these technologies, these requirements are out of reach for many researchers and citizen scientists. There are many important scenarios where analyzing these data without high-end computing hardware is desirable, especially in remote environments. Interestingly, current iOS devices, including both iPads and iPhones, have significant computing resources, with clock speeds and onboard RAM approaching that of high-end laptop computers. That said, no standalone genomics analysis software is currently available for iOS devices.

Addressing this critical gap, we have developed iGenomics, an iOS application that allows anyone to easily align and analyze DNA sequences in a mobile environment. iGenomics utilizes the same high performance algorithms for read alignment and variant calling as mainstream software, although iGenomics marks the first time these algorithms have been implemented in a mobile iOS environment. Additionally, using the advanced user interface features available in iOS, iGenomics allows for interactive visualization and inspection of the read alignments and variant calls, and contains additional features for reviewing critical mutations of interest. For example, iGenomics comes bundled with a listing of critical mutations in the influenza A virus that indicate which antivirals are most likely to be ineffective (Hussain et al. 2017).

Due to the lower amount of processing power in mobile devices compared to high-end desktop computers or servers, iGenomics is limited in the size of the genome that can be processed. However, the implementations in iGenomics have been rigorously tested through direct comparisons with the BWA-MEM/Samtools framework for alignment and variant calling for viral and microbial genomes. All alignment and analysis algorithms employed by iGenomics have been tested on both real and simulated datasets to ensure consistent speed, accuracy, and reliability of both alignments and variant calls. Consequently, iGenomics is leading the shift of DNA analysis software and sequencing tools towards mobile devices and marks a great leap forward towards widespread DNA analysis by non-bioinformatician doctors, researchers and citizen scientists. Furthermore, iGenomics is available open-source to facilitate mobile genomics technology research and, in turn, accelerate the speed at which this technology is developed.

## Results

### Interactive Sequence Analysis on your Smartphone

iGenomics brings a high level of interaction to DNA sequence analysis (**Figure 1**). Common touchscreen gestures allow for users to browse the alignment data in an easy-to-use and intuitive manner. This allows for the app to be used with almost no learning curve.

**Figure 1:**
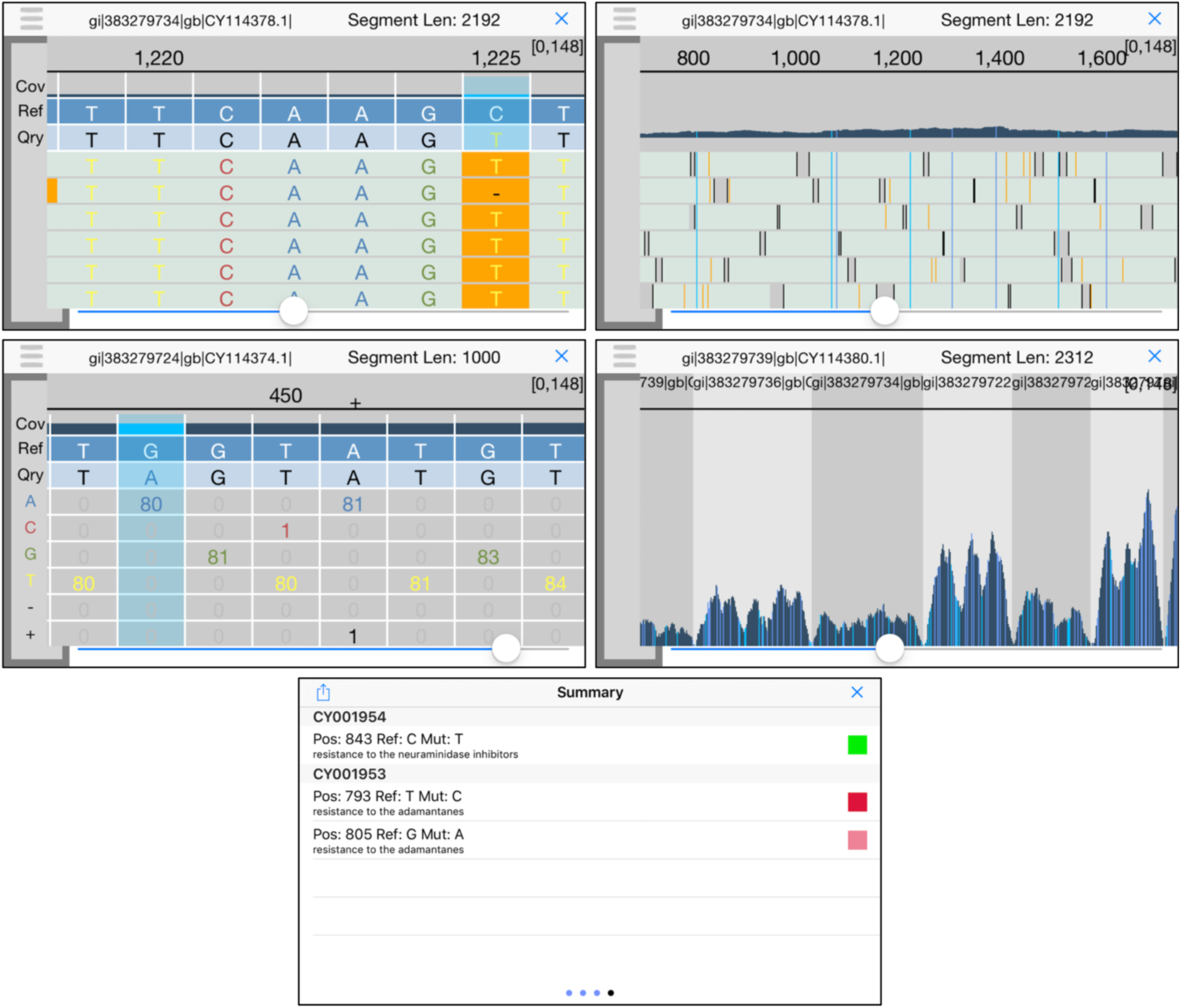
iGenomics iPhone screenshots **(top-left)** Alignments display; **(top-right)** Alignment display zoomed-out; **(middle-left)** Coverage profile; **(middle-right)** Coverage profile zoomed-out, **(bottom)** Known mutations display. In the known mutations display, green indicates the mutation is not present, dark red indicates the listed mutation is present and the mutation is homozygous, and pink indicates the listed mutation is present and the mutation is heterozygous. In both the alignments display and coverage profile, there is an indicator in the top right of the form [X, Y] that represents the minimum coverage X across all positions and maximum coverage Y across all positions.

The first step of analysis is selecting the reads and a reference genome for analysis in either fasta or fastq format. iGenomics provides multiple options for inputting both reads and reference files: selecting from a variety of default files for common bacterial genomes, using Dropbox to choose a file, or loading a fasta or fastq file straight into iGenomics from another app (such as Google Drive) or Airdrop. Then, from a single view, the user can choose the reads file, the reference file, and, optionally, a tab-delimited file annotating known important mutations. For example, iGenomics comes with a preloaded known mutations file that indicates certain mutations in the influenza genome, which, if present, cause resistance to certain antivirals (Hussain et al. 2017). This single view design is meant to be simplistic and requires minimal user effort. After choosing the files to align, the user can either select the “Analyze” button to align reads to the reference genome using the default parameters or can choose to configure certain parameters before aligning. The parameters available include the maximum error rate for alignments and to enable trimming for fastq files.

After aligning completes, the user is brought to the analysis pane. The main view, known as the alignments display, is an IGV-like rendering of how the reads are aligned to a reference genome, with the ability to scroll left, right, up, and down through all of the aligned reads. Aligned bases that differ from the reference base are highlighted in a different color, as are consensus calls. A long-touch on a read presents additional details about the read, including the read name, the edit distance of the alignment, the gapped read and gapped substring of the reference genome the read aligned to, and whether the forward read or the reverse complement aligned. The user can also use the pinch-gesture to zoom out, revealing a high-level overview of the individual alignments as well as a coverage profile of the number of reads that aligned at each position. Mutations are still highlighted after zooming out, allowing the user to see where all of the mutations occur in one view.

Another powerful view within the analysis pane is the coverage profile, which displays the count of each base that aligned at each position. Positions where the reference base does not match the base of the reads are highlighted so that the user can see that this position contains a mutation (heterozygous mutation are highlighted with a different color). To scroll through the coverage profile, the user simply has to swipe left or right. If a user would like to view more detailed information about a given position, he/she simply holds down any of the boxes in that position and an informative view elaborating upon the position’s contents will pop up. By using the pinch gesture to zoom-out, the user reveals a graph of the number of reads that aligned at each position, resembling that of the zoomed-out alignments display but with a full-screen graph.

The Summary window, accessible from within the analysis pane, has four pages and provides some useful tools for high-level analysis. The first page provides buttons to view the alignments display, coverage profile, coverage histogram, and list of all found mutations. The coverage histogram graphs the frequency of each level of coverage, specifically the frequency of a particular number of reads aligned to a position, and is overlaid by a Poisson curve for context. Within the list of all found mutations, the user can scroll through all mutations, and then select one to inspect that position in the analysis pane. The second page gives an overview of the alignments, including the percent of reads matched, the total number of reads input, the number of mutations, and the names of the reads and reference files. This page also provides the user with the capability to search for positions in the reference genome by position or by a query string, which uses BWT exact match for rapid searching. The third page contains a large picker view that allows the user to intuitively move between sequences/segments in the reference genome. The last page contains a list of known mutations if the user selected a known mutations file during the file input stage. This list contains mutation position, mutation details (such as resistance to antivirals), and a color-coded indicator denoting if a mutation was found at that position and if that mutation indicates a known mutation.

### Simulated read runtime analysis

In order to observe the efficiency and accuracy of iGenomics running on an iPhone 8, we first tested several simulated data sets. The reference genomes we used were:

1. phiX174, a widely used control sequence for Illumina sequencing (Genbank:NC_001422.1);
2. a Zika virus genome (isolate Zika virus/H.sapiens-tc/KHM/2010/FSS13025);
3. a H3N2 influenza genome (A/California/7/2004(H3N2));
4. a H1N1 influenza genome (A/New York/205/2001(H1N1)); and
5. an Ebola genome (isolate Ebola virus/H.sapiens-wt/SLE/2014/Makona-G3686.1).

From these reference genomes, we then simulated reads using DWGSIM (https://github.com/nh13/DWGSIM) according to the following conditions: the average coverage is 100x, the genetic mutation rate was set to 0.5% and the read characteristics would mirror reads produced by real-world sequencers. Accordingly, reads of length 100bp and sequence error rate of 1.0% were simulated to mirror reads generated by Illumina sequencers and reads of length 1,000bp and sequence error rate of 10.0% were simulated to mirror reads generated by Oxford Nanopore sequencers. For comparison purposes, we also measured the runtime when aligning and identifying variations using a BWA-MEM (Li 2013) using “-x ont2d” and Samtools pipeline for the same datasets. Notably, iGenomics uses an FM-index and banded dynamic programming implementation similar to BWA-MEM allowing the analysis to focus on major differences in hardware.

When comparing the runtime of iGenomics against datasets with different genome lengths, we observe a nearly linear relationship between genome length and alignment runtime (**Figure 2**). This is explained by a powerful feature of the BWT in which the time for an alignment of a single read is essentially independent of genome size. Consequently, since the simulations use a consistent amount of coverage per genome, the linear increase in runtime is explained by the linear increase in the number of reads to align. It is also worth noting that the iGenomics trend-lines closely follow the pattern of those of BWA-MEM+Samtools. This both adds credibility to iGenomics as a sequence alignment and analysis tool and to the field of portable genomics, as all of these important viruses can be analyzed in under 5 seconds on a mobile device.

**Figure 2:**
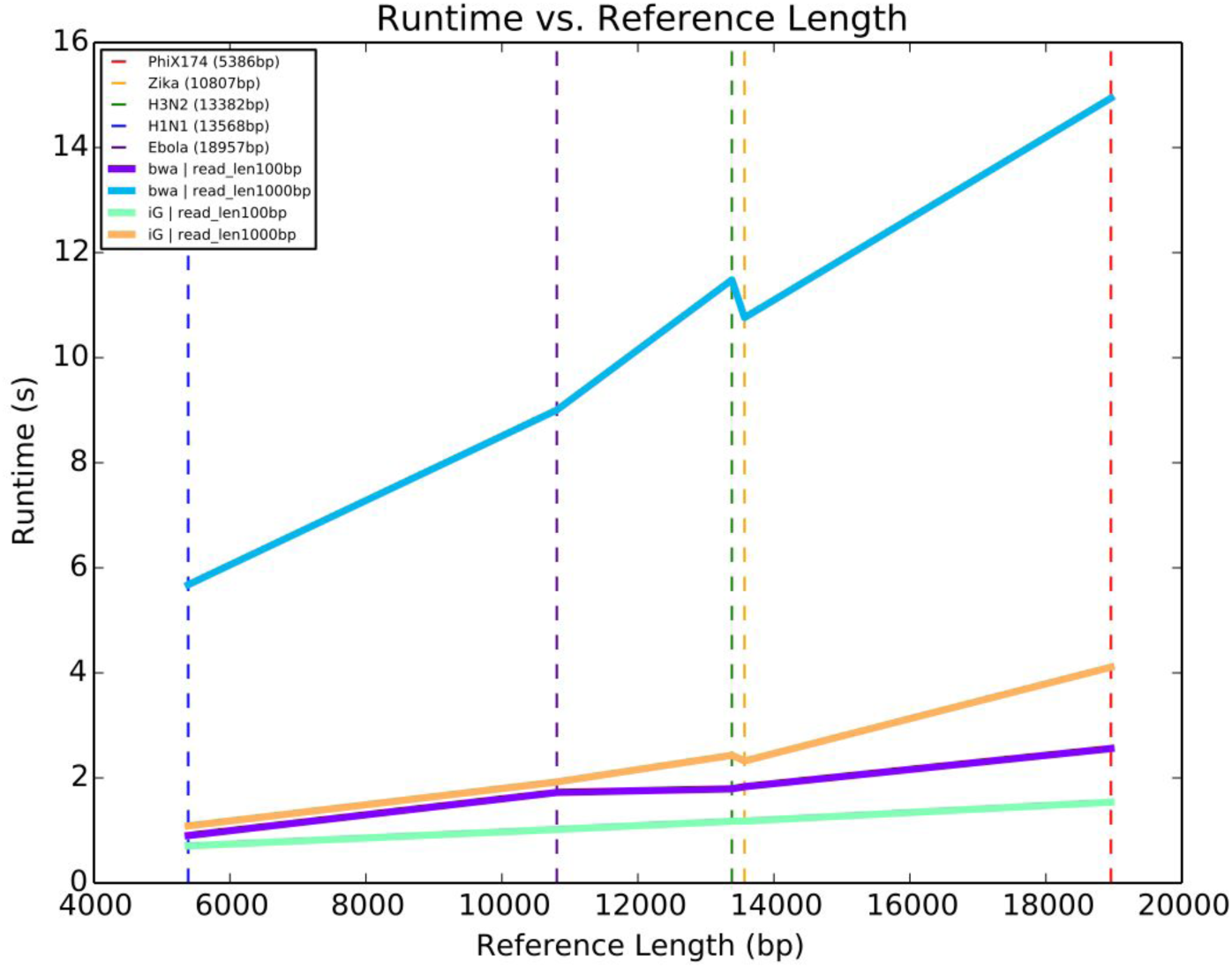
Runtimes for simulated reads from five reference genomes. The data sets consisted of reads averaging 100x coverage and a reference file. Each data set was tested, defined as aligning then variant calling, using iGenomics and a BWA/Samtools pipeline. Each trend line indicates the runtime for each data set using the denoted alignment and analysis software-iG for iGenomics and bwa for the BWA/Samtools pipeline. The dotted lines indicate the specific measurements recorded.

### Simulated read accuracy analysis

We next evaluated the accuracy of iGenomics using reads simulated from the H1N1 Influenza genome (same sample as above). In each trial, we simulated an average of 100x coverage for all combinations of the following sets of parameters: sequence error rates of 0.01, 0.1, and 0.2, mutation rates of 0.001, 0.01, and 0.1, and read lengths of 100bp, 250bp, and 1,000bp. The range of the simulation parameters is designed to test iGenomics across a variety of different possible sets of reads that iGenomics could be used with. After simulating the read sets, each simulated sample was independently aligned to an H1N1 reference genome using iGenomics. For each sample, we recorded the runtime and the reported list of mutations found. In order to check the validity of the mutations found by iGenomics, the reported mutations were compared to the DWGSIM-generated list of simulated mutations. We then compare the variants reported by iGenomics to DWGSIM, allowing for up to 5bp differences to account for ambiguity that can occur, especially indels within locally repetitive sequencing. Key metrics that were evaluated relative to DWGSIM were precision, recall, and F-Score (the harmonic mean of precision and recall).

The results of the comparisons between iGenomics’ reported mutations and DWGSIM’s list of mutations confirm iGenomics accuracy. Most datasets show a high-degree of accuracy (F1) well over 90% (**Figure 3**). The few experiments with lower precision or recall occur with the most difficult scenarios of the highest sequencing error rate and the lowest mutation rate. For comparison, the same results were also computed with input from a BWA-MEM/Samtools pipeline. Interestingly, iGenomics tends to exhibit a higher degree of recall, precision, and overall accuracy.

**Figure 3:**
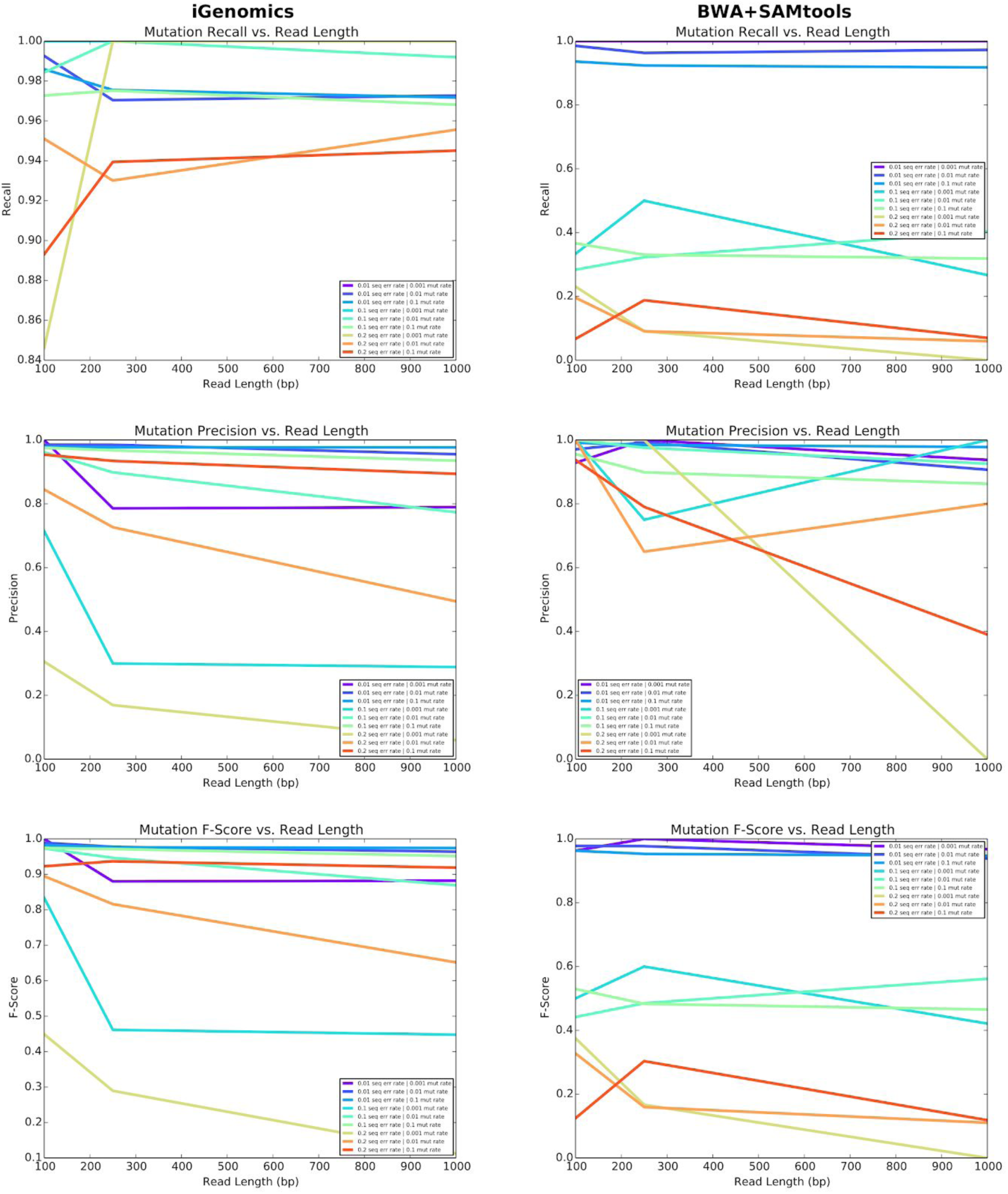
Mutation identification accuracy for simulated H1N1 flu datasets of varying mutation rates and error rates for iGenomics (left) and the BWA-MEM/Samtools (right) pipeline. The top, middle, and bottom plots show recall, precision, and F-score, respectively.

Another important consideration for iGenomics is the runtime required. The runtime of iGenomics for each of these simulated data-sets was below 3 seconds (**Figure 2**). Furthermore, iGenomics aligned reads and identified mutations in these simulated datasets about 4x to 5x faster than the BWA-MEM/Samtools pipeline (**Figure 4**). For context, the BWA-MEM/Samtools runtime for these data sets was computed on an early 2015 MacBook Pro with a 2.9GHz Intel Core i5 running OS X El Capitan while the iGenomics runtime was computed on a 2017 iPhone 8 with a 2.39 GHz A11 Bionic Chip running iOS 12.3.1.

**Figure 4:**
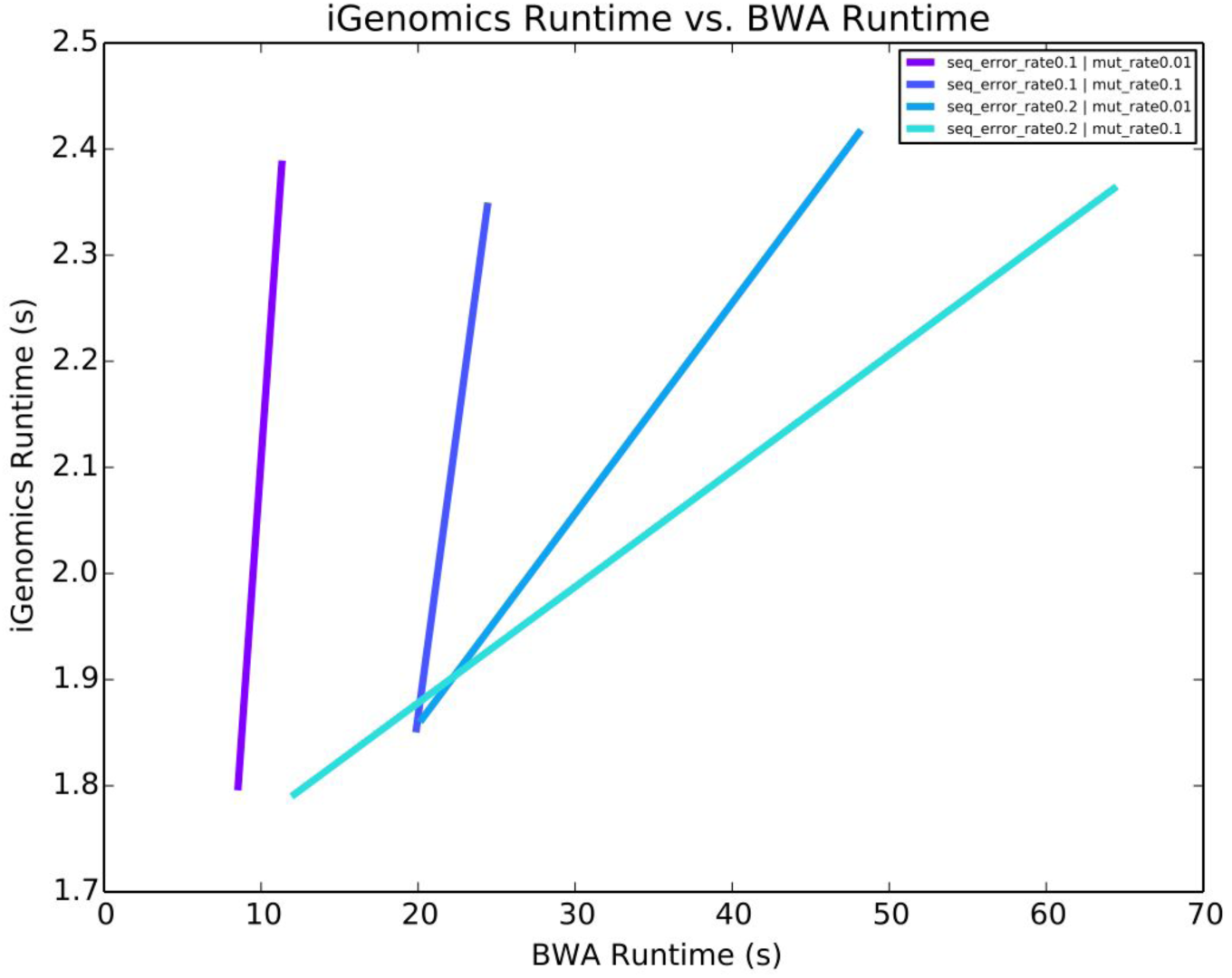
iGenomics runtime vs. BWA/Samtools pipeline runtime for simulated datasets of varying mutation rates and sequence error rates of H1N1.

### Viral Genome Analysis

iGenomics was next tested on several clinical and environmental viral samples sequenced using the Oxford Nanopore MinION in order to demonstrate both the functionality and accuracy of iGenomics relative to standard tools such as BWA-MEM and Samtools. The purpose of these tests is to show the overall utility of iGenomics as a mobile counterpart to desktop aligners and analysis software typically used by researchers and as a novel sequence analysis platform.

These tests focused on public MinION data from Ebola (sample https://raw.githubusercontent.com/nickloman/ebov/master/data/fastq/004674.2D.fastq from (Quick et al. 2016)), and Zika (sample http://s3.climb.ac.uk/nanopore/primal_KX369547_R9.tgz from (Faria et al. 2016)), as well as MinION and MiSeq data from a clinical H3N2 sample we previously collected (A/New York/A39/2015 (H3N2)) (Ding et al. 2019) (**Methods**). The Ebola trial focused on comparing iGenomics found mutations to those found by Samtools using the isolate Ebola virus/H.sapiens-wt/SLE/2014/Makona-G3686.1 as the reference (GenBank: KM034562.1). For Zika, the test was based on using a ground-truth set of mutations derived from a consensus genome using the isolate Zika virus/H.sapiens-tc/KHM/2010/FSS13025 (GenBank: KU955593.1) as the reference. The H3N2 test was designed to demonstrate iGenomics consistency across data produced by different sequencers by comparing the results of the Nanopore and MiSeq data when aligning to the isolate (A/California/7/2004(H3N2)) genome.

In all of the cases examined, iGenomics had a faster runtime than the desktop alignment pipeline of BWA-MEM/Samtools (**Figure 5**). This is likely due to a difference in how iGenomics and the desktop software store the alignments in memory. Since iGenomics is targeted to be a focused mobile analysis platform for small genomes, iGenomics needs to run very rapidly. Instead of separately reporting each alignment and writing the alignments to disk, then separately sorting the alignments, and then scanning for variations, as BWA-MEM/Samtools does, iGenomics records the full gapped alignments and coverage profile matrix in RAM so that the subsequent mutation identification can avoid repeating computations. Furthermore, iGenomics keeps this data in RAM until the user exits the analysis screen to allow for exploring the various visualizations and performing interactive analysis with negligible lag time. This presents a standard time vs RAM tradeoff present in many software applications, and here we have elected for fast processing to ensure the application is as responsive as possible.

**Figure 5:**
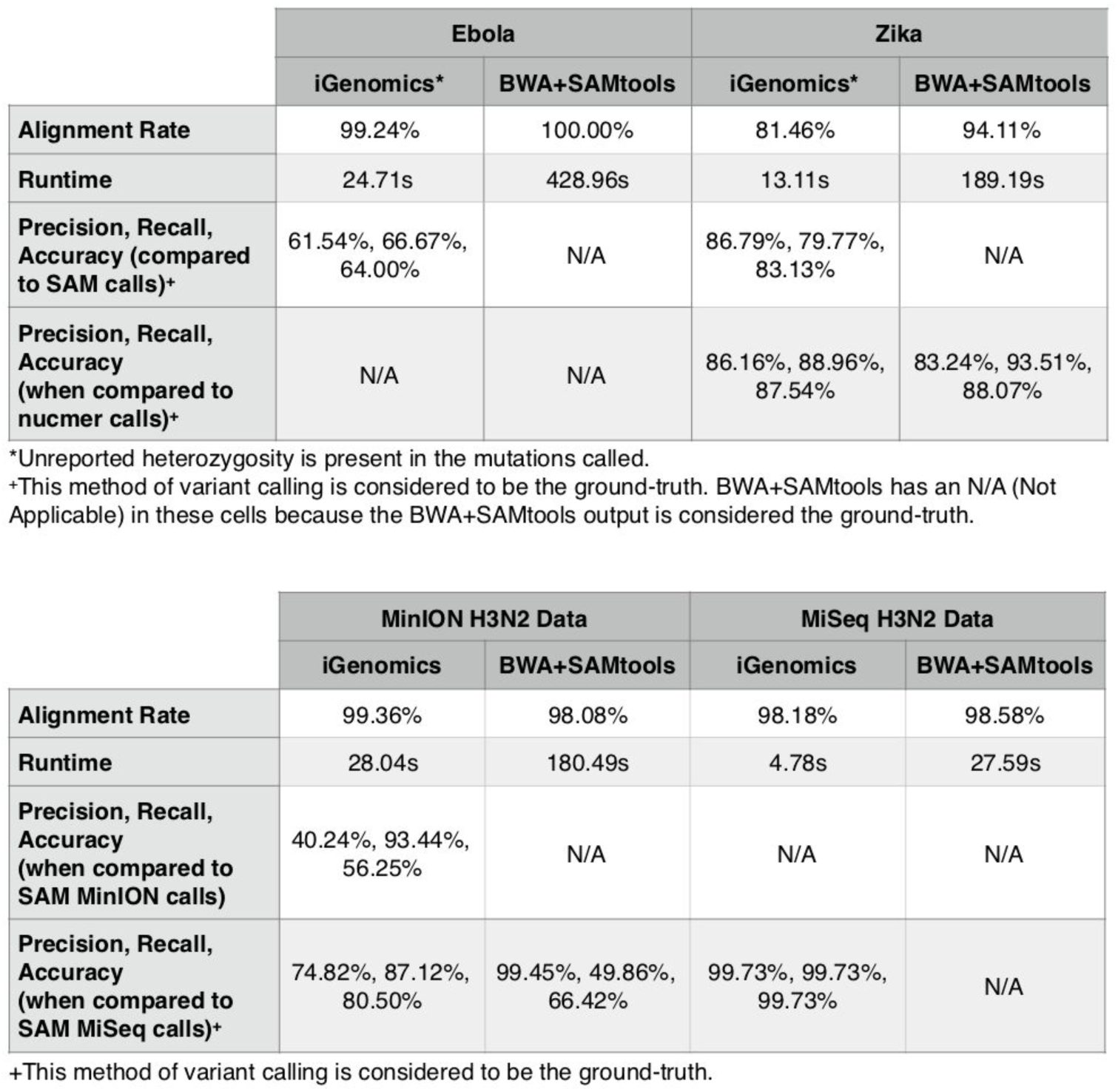
Comparison between iGenomics and BWA-MEM/Samtools pipeline for real reference genomes and reads obtained from MinION (Nanopore) and MiSeq sequencers.

### Influenza typing

Influenza disease is caused by RNA viruses from the family Orthomyxoviridae (Krammer et al. 2018). There are three distinct viral types, A, B, and C that can infect humans. Influenza types A and B cause the annual epidemics, while influenza C is generally less severe. The influenza A genome is organized into eight segments, and is classified into subtypes based on genetic variants within the two proteins on the surface of the virus: hemagglutinin (HA) and neuraminidase (NA). There are 18 different hemagglutinin subtypes and 11 different neuraminidase subtypes (H1 through H18 and N1 through N11, respectively). Many of the major influenza pandemics have been caused by influenza type A infections. For example, the 1918 flu pandemic (the “Spanish flu”), was caused by a deadly Influenza A virus strain of subtype H1N1, and the Hong Kong Flu in 1968 was caused by the H3N2 subtype. Consequently, the type and subtype of an unknown influenza sample is extremely important and urgent to determine.

As a final demonstration of how iGenomics can be used, we also considered an influenza identification task where influenza sequencing data are aligned to several strains of flu at the same time in an attempt to determine the type and subtype. For this, we developed an influenza “pan-genome reference sequence” containing representatives for three different Influenza genomes related to antigenic strains that were circulating from 2009 to 2016: H1N1pdm09 (A/California/04/2009), H3N2 (A/Brisbane/10/2007; A/Perth/16/2009; A/Texas/50/2012; A/Victoria/361/2011; and A/NewYork/03/2015), and Influenza B (B/New York/1352/2012). For this analysis, segments that are shared across influenza A subtypes were only reported once. For the pan-genome, we also include a catalog of mutations in these genomes that have specific variants known to reduce the efficacy of antiviral treatments. The identity of the A segment is identified by evaluating which of the potential segment types has the largest number of alignments.

In order to test alignments against the pan-genome, we ran iGenomics using simulated MinION (1,000bp, sequence error rate 10.0%) and Illumina (100bp, sequence error rate 1.0%) reads from pH1N1 and H3N2 with mutations rates 0, 0.001, and 0.005. After alignment, we evaluated if the reads were correctly aligned to the type and subtype that they originated from. If the alignment matches the segment of origin, we consider that alignment “passing”. The segment identification rate is the number of passing alignments divided by the total number of alignments. The results of this experiment show that we have a greater than 93% identification rate, meaning that in most cases this simple process can accurately and quickly determine the type and subtype of the flu genome entirely on a mobile device (**Figure 6**).

**Figure 6:**
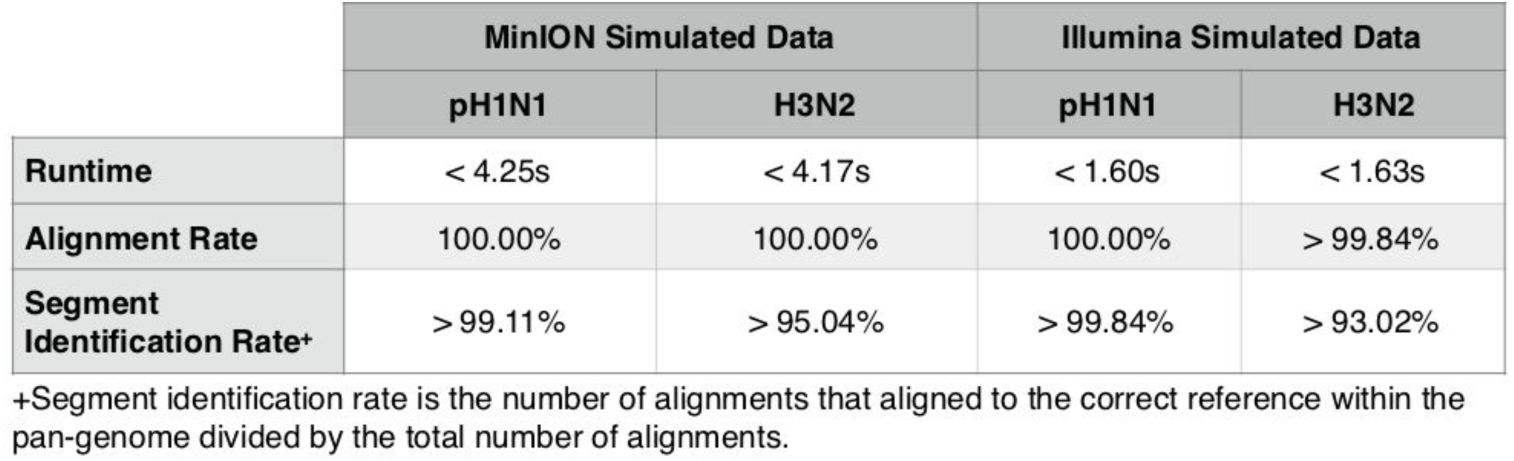
Table indicating alignment details for simulated datasets aligned using iGenomics to a pan-genome composed of multiple Influenza genomes. The pH1N1 reads were simulated from the H1N1pdm09 (A/California/04/2009) genome and the H3N2 reads were simulated from the H3N2 (A/NewYork/03/2015) genome.

## Discussion

DNA sequencing has advanced tremendously over the past three decades; a process that once required large million dollar instruments can now be done on handheld devices costing only $1,000. However, it is important to consider that sequenced DNA reads themselves provide little information without software to align and analyze them. For high-end servers and laptops, this software already exists; for mobile devices, iGenomics is the first comprehensive solution for researchers and citizen scientists to easily analyze sequence data using a device that they already own.

Unlike traditional DNA mapping software, iGenomics can be used in virtually any location because of the inherent portability of mobile devices like the iPad and iPhone. iGenomics implements the same advanced bioinformatics algorithms that are used for rapid alignment and analysis for other platforms. Consequently, the true novelty of this application is not in the algorithms used, but rather how they have been implemented in a mobile environment. The entire workflow for iGenomics is designed to be very simple and intuitive. A user effortlessly picks a reads file to analyze and, once selected, the alignment, variant calling, and visualization are completed within seconds. This is accomplished without any internet connectivity through an optimized implementation in Objective-C. Interestingly, while Objective-C is sometimes an afterthought for computationally intensive apps, iGenomics leverages the language’s capabilities to generate both a unique user experience and fast analysis times.

iGenomics is designed for quickly computing detailed genetic information about specific mutations within different viral or bacterial genomes. A practical use case of iGenomics could be a researcher with limited computational resources sequencing cDNA of an Influenza sample, loading and aligning the cDNA reads with iGenomics, and getting a first analysis of the Influenza’s treatability. Remarkably, in just a few seconds, the user would be able to make a first analysis of which antivirals the input strain is resistant to. A promising capability of iGenomics is its ability to load reference genomes and reads from outside sources, perform alignment and variant calling, and export the results all without any internet access. For example, by using Airdrop to both import and export data from iGenomics, a researcher can analyze DNA in remote locations without any internet connectivity.

Future developments for iGenomics are far reaching as DNA sequencing instruments continue to evolve to the point where they could be directly attached or integrated with mobile devices. In fact, Oxford Nanopore has announced that they hope to have a new sequencer, named the “SmidgION”, that connects directly to iOS devices available for researchers within the next year. At that point, using mobile sequencing technology with iGenomics, DNA can truly be sequenced, aligned, and analyzed anywhere and absolute mobility of the genomics field will be achieved. As the processing power and memory contained within mobile devices improves, so will the overall performance of iGenomics in handling even larger and more complex samples.

## Methods

The implementation of iGenomics follows the state-of-the-art algorithms and data structures used in standard bioinformatics applications. However, the visualization of the read alignments and mutations is unique to iGenomics and was created with the intention of allowing the user to have powerful analysis capabilities while still maintaining a simplistic mobile-friendly interface.

### 1. Indexing the genome with the Burrows-Wheeler Transform (BWT)

The Burrows-Wheeler Transform (BWT) is constructed by lexicographically sorting the cyclic permutations of the input genome appended by a end-of-string character. By convention, we use a dollar sign (‘$’) as the end-of-string character, which has a lexicographical value less than any letter in the English alphabet and ensures the end of the original sequence can be found. This sorted list creates what is known as the Burrows-Wheeler Matrix (BWM). Then, to extract the BWT from the sorted permutations, the last character of each row in the matrix is extracted in order and appended to a string.

To first lexicographically sort the cyclic permutations, a quick and efficient sorting algorithm must be used so that this function is fully optimized. iGenomics uses a version of QuickSort, a divide-and-conquer sorting algorithm, because on average it takes O(n log n) time for n objects to be sorted. Finally, to obtain the BWT from the sorted array, the final character of each row in the matrix is copied into a string with the first character copied having the first position, the second character copied having the second position, and so forth.

**Figure 7:**
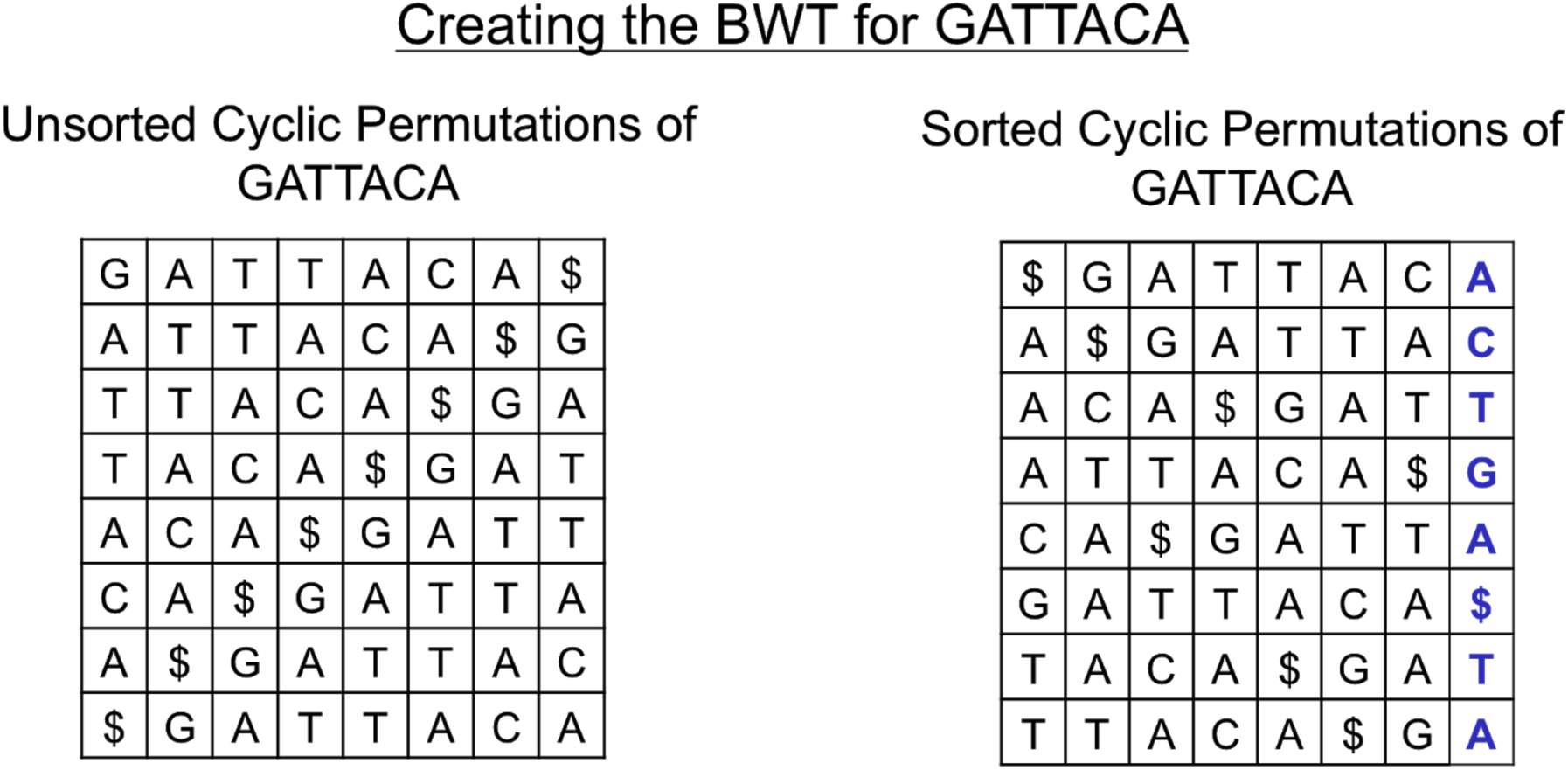
Diagram of how the Burrows-Wheeler Transform is created. (left) All cyclic permutations of the text “GATTACA”. (right) The Burrows-Wheeler Matrix of the text consisting of the sorted cyclic permutations of the text.

### 2. Read alignment

iGenomics uses a seed-and-extend process for read alignment in which first relatively short exact matches, known as seeds, are found using the BWT, after which they are then extended into end-to-end alignments using dynamic programming. The seed size is based upon the maximum edit distance (a user-specified parameter) allowed for a read that successfully aligns to be considered a match. The maximum edit distance is inputted as a decimal value edit rate, and multiplying that value by the length of the given read will give the maximum possible edit distance we allow when aligning that read. During the aligning process, each read is split into the edit distance plus one segment of equal length. This relies on the widely used technique that if the string matches with at most X edits, then at least 1/(X+1) of the segments must still match without error (Baeza-Yates and Perleberg 1996). For example, if the user allows only 1 edit, the algorithm divides the read into left and right halves (1/(1+1)) knowing that the correct alignment will include an exact match of one of those segments.

Exact matching means finding all of the places in the reference genome where a given query matches exactly, character-for-character across its entire length (Langmead, 2012). To do this effectively, the trait of the BWT known as the Last-First Property is used as the basis for an exact matching algorithm. The Last-First property states that the occurrence of any character in the last column of the BWM, which is the BWT, corresponds to the same occurrence of that character in the first column of the BWM. Using the first column of the BWM and the BWT to create an FM-index, the algorithm navigates the rows of the index which contain exact matches and then converts these positions from the BWT to positions in the reference genome.

**Figure 8:**
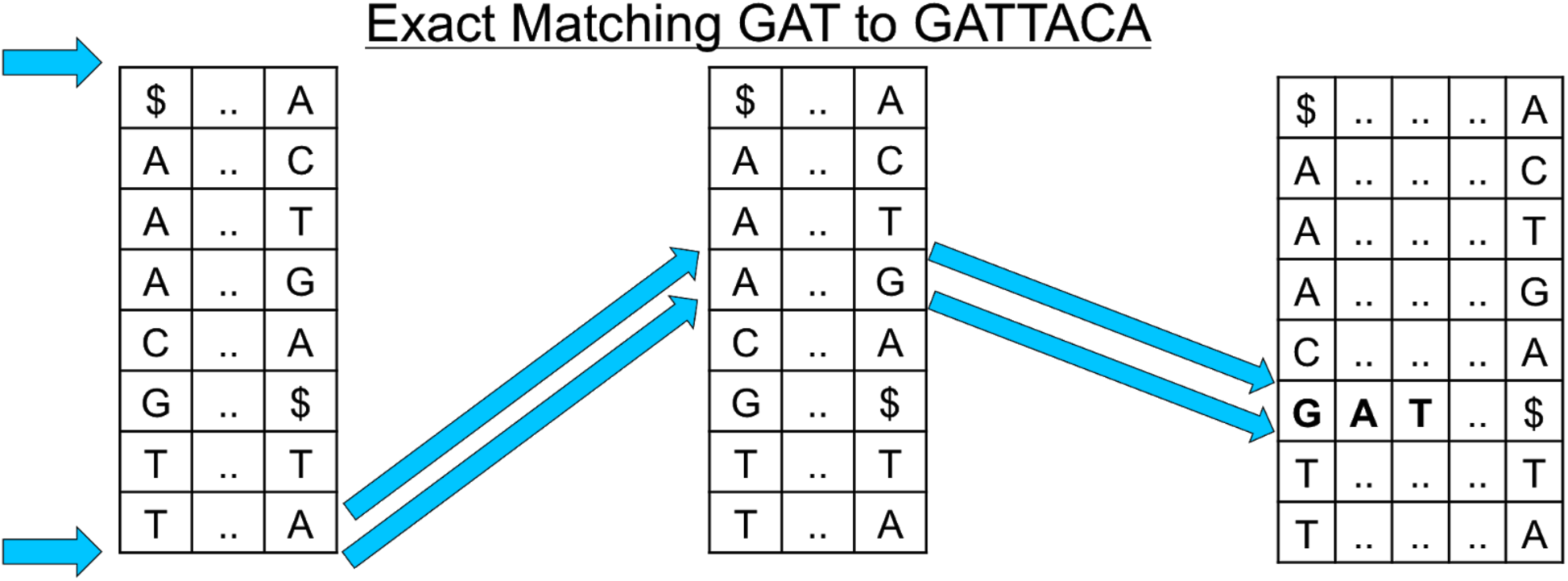
A diagram showing the exact match algorithm by repeated application of the Last-First property using the characters of the query string.

After the seeds are found, iGenomics computes the end-to-end edit distance allowing for substitutions as well as insertions and deletions (Smith and Waterman 1981). To make this as efficient as possible, iGenomics uses a banded computation. This method works by only computing a subset of the dynamic programming matrix, a band of the edit distance table, with the band having a standard width of (the maximum edit distance * 2 + 1). To determine where to begin the band computation, iGenomics attempts to exact match a 20bp substring of the read. If the exact match is successful, the banded distance will be computed relative to the matched position of the substring. If the exact match is unsuccessful, an exact match with the 20bp substring of the read starting at the second character will be attempted. This process continues with the substrings continuously moving one character over until either the read successfully aligns or none of the exact matched 20bp substrings yields a successful alignment.

**Figure 9:**
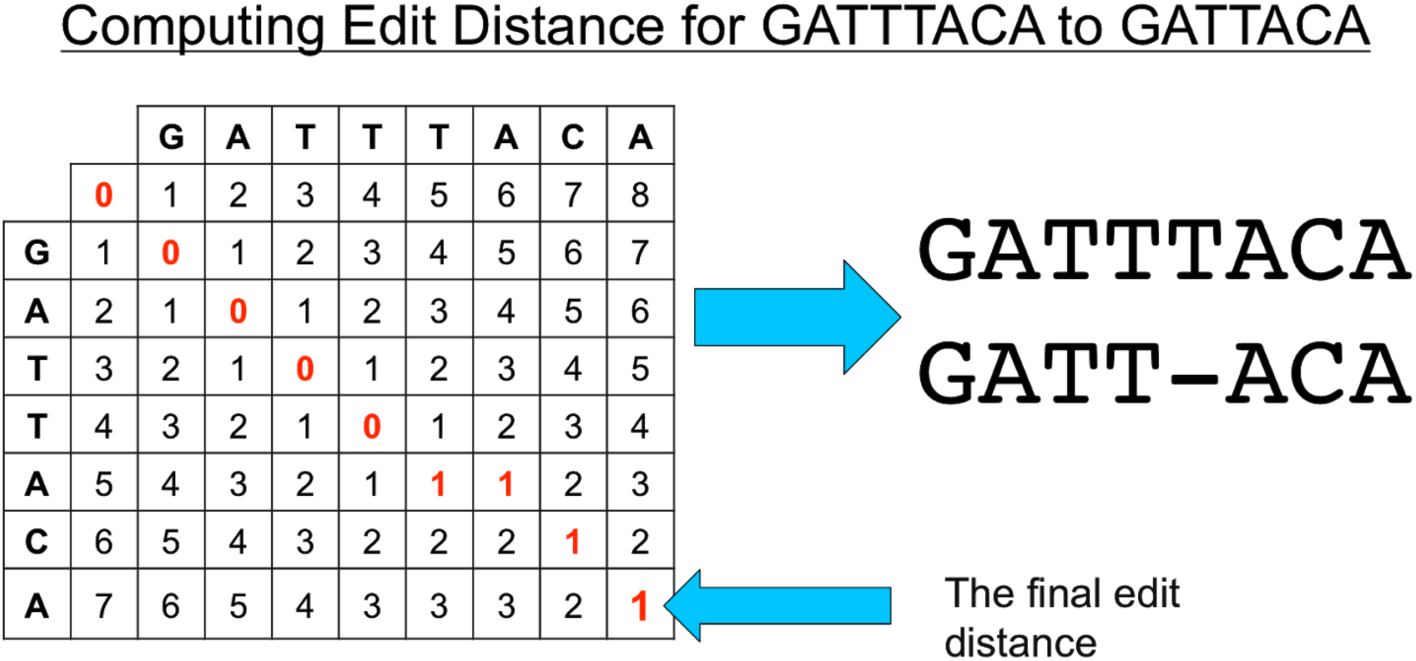
A diagram showing how edit distance is computed for two strings. Each cell of the matrix represents the minimum of three possible values: 1) the left cell plus one (representing the cost of adding a gap on the left string); 2) the upper cell plus one (representing the cost of adding a gap on the top string; and 3) the upper left cell plus zero, if the top string equals the left string, or one, if the characters do not match to account for the cost of another substitution.

### 3. Coverage profile and variant identification

The coverage profile concisely summarizes how the reads are aligned to the genome. The internal data structure for the profile is a coverage profile matrix, which spans the genome and at each position contains a row for the number of: matched base-pairs, A, C, G, T, and (non-base-pair) deletion characters. The matched positions of each read are tallied and the characters of the read are added, so that the positions of the matrix that the read overlaps are marked within the matrix. Once the coverage profile matrix is completely generated, variants can be identified, a graphical representation of the profile can be formed, and the number of alignments can easily be seen.

**Figure 10:**
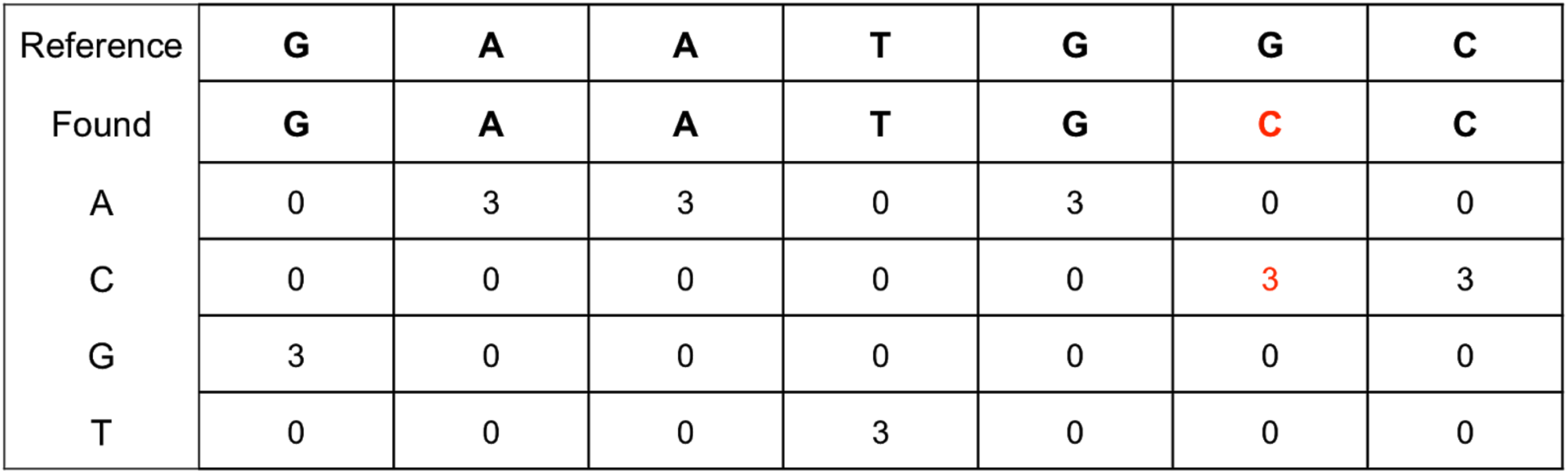
A table showing how the coverage profile is represented within iGenomics, summarizing how the reads align to the reference genome (an example of reads aligned to a reference genome is shown in Figure 1).

Variants are identified by scanning the array of matched characters, and at each position if the matched character differs from the reference character, a mutation, or variant, would be reported (Li et al. 2009). The major challenge of this analysis is distinguishing sequencing errors from real mutations, and differentiating between homozygous and heterozygous mutations. In a diploid genome, homozygous mutations are mutations that occur on both copies of a chromosome whereas heterozygous mutations occur on one copy of a chromosome but not both. iGenomics recognizes heterozygous mutations as positions in the genome where there is a nearly equal coverage of more than one base existing in the set of aligned reads according to a user-specified relative minimum heterozygosity threshold. Thus, if two or more bases at a position have relative coverages greater than that threshold, the mutation present at that position is considered to be heterozygous. In haploid species, such as the viral and bacterial pathogens described above, this threshold is used to find variants that occur within a minimum allele frequency within the population.

Immediately after alignment has completed, each position within the reference genome is assigned a value indicating whether the reads at that position matched either exactly, heterozygously, homozygously, heterozygously where there is a known mutation, or homozygously where there is a known mutation. This allows iGenomics to highlight all mutations with their associated heterozygosity and importance. Known mutations are loaded through a user-inputted text file. This file contains each known (important) mutation’s reference base, mutated base, position, segment (or chromosome) the mutation is expected to occur in, and a free-text description of what this mutation indicates. The known mutations functionality enables iGenomics to be specifically targeted for the analysis and treatment of different genomes, such as known mutations associated with Influenza antiviral resistance.

### 4. Visualizations and interactive analysis

The main challenge with the GUI was to create one that was both useful and unique when compared to other desktop DNA analysis software. The key to achieving these goals was to take advantage of the distinctive features of the iOS environment. Ultimately, a custom graphics engine was built to handle the constant redrawing of the analysis interface and, visually, this engine sits on top of Apple’s CoreGraphics library. In addition to the analysis interface, a utility interface was developed, which contains features for rapidly analyzing and quickly navigating the alignments.

The solution to developing this unique interactive analysis screen was to employ many touch-related functions that are natural to anyone who has ever used a touch screen mobile device. Scrolling requires a simple finger drag while viewing a large-scale version of the coverage profile merely requires performing a pinch gesture on the screen. The information pertaining to mutations can be viewed at any position by tapping on one of the reference genomes or found genome boxes at that position. Even this action takes advantage of the mobile iOS environment because a popover view is used to display the information at the tapped position. At the bottom of the screen, there is a variable scrubbing speed slider so that the user can move across the genome quickly or at a slower rate by dragging up while moving the slider.

Simple functions such as searching for a specific query or position are also included in the analysis view. To minimize clutter on the screen, when a user searches for a certain string, he/she is instantly taken to the next occurrence of that string, as opposed to displaying a large list of positions to the user. One of the most notable of these functions is the ability to change the minimum relative heterozygosity value (known as mutation coverage within iGenomics) on the fly through a slider. Once the user has concluded analyzing on the mobile device, he/she has the option to export mutations and analysis data via a variety of means: email, Dropbox, Airdrop, or sharing via installed apps (such as Google Drive). The mutations are outputted in a VCF (Variant Call Format) file format so that they are compatible with traditional desktop analysis software.

## Flu Isolate Sequencing

### Sample collection and amplification

Clinical specimens of nasopharyngeal swabs were collected from patients in New York City in the 2014-2015 flu season as previously described (Ding et al. 2019). The specimen used in this study was designated as A/New York/A39/2015 (H3N2) and is available in the SRA as sample ID SAMN08454624. Briefly, the RNA was eluted in 30 µl of RNase-free water and 3 µl was used as a template for the amplification of the entire influenza A or B genome using previously described Multi-segment RT-PCR (M-RTPCR) method (Zhou et al. 2009). The presence of the cDNA copies of the genomic segments were examined by running 3 µl of the M-RTPCR amplicons on a 0.8% agarose electrophoresis gel. The influenza genomic amplicons were purified using a 1x Agencourt AMPure XP purification step and assessed by Qubit analysis to quantify the mass of the double-stranded cDNA present.

### Nanopore MinION sequencing

The library preparation and sequencing procedures were performed following manufacturer’s instructions for the Nanopore Sequencing using the SQK-MAP006 kit. Purified DNA was used for end repair and dA-tailing, followed by 1x AMPure XP beads purification. The resultant DNA was quantitated by Qubit analysis and the molarity was further determined by using Agilent 2200 TapeStation system with a Genomic DNA ScreenTape. Next, 0.2 pmoles of the DNA was used in adaptor ligation, and the reaction was purified using MyOne C1-beads. The final DNA was eluted in 25 µl Elution Buffer and is called Pre-sequencing Mix. For the SQK-MAP006 sequencing kit, 12 µl Pre-sequencing Mix was combined with 75 μl 2x Running Buffer, 59 μl nuclease-free water, and 4 μl Fuel Mix and then loaded into the FLO-MAP003 flow cell. A re-loading was also performed. The sequencing was run on the MIN-MAP001 MinION sequencing device, which was control by the MinKNOW software using the MAP_48Hr_Sequencing_Run.py script provided by Oxford Nanopore or using the MAP_140to5xVoltage_Tuned_plus_Yield_Sequencing_Run.py script provided by John Tyson. Raw data was uploaded to the cloud-based Metrichor platform and basecalling was performed using the application of 2D Basecalling for SQK-MAP005 Rev 1.62 or 2D Basecalling for SQK-MAP006 Rev 1.62.

### Illumina MiSeq sequencing

The sample was prepared for sequencing on the Illumina MiSeq platform according to the manufacturer’s protocol (15039740 v01) as previously described (Ding et al. 2019). Sequencing data was then generated by a 2×300bp run using an Illumina MiSeq 600 Cycle v3 reagent kit.

## Data Availability

All sequencing data (genuine and simulated) along with a tutorial on iGenomics are available online: http://schatz-lab.org/iGenomics/.

## Acknowledgements

We would like to thank Jaak Raudsepp for his helpful discussions and involvement during the development of iGenomics. We would also like to thank Dr. Mirella Salvatore for providing the flu samples. The project was supported in part by National Science Foundation award (DBI-1350041) to MCS. This paper is dedicated to Albert Palatnick, grandfather of Aspyn Palatnick, who has fueled Aspyn’s fascination with science, technology, and innovation since he was a child.

