## Supplemental Figures for "iGenomics: Comprehensive DNA Sequence Analysis on your Smartphone"

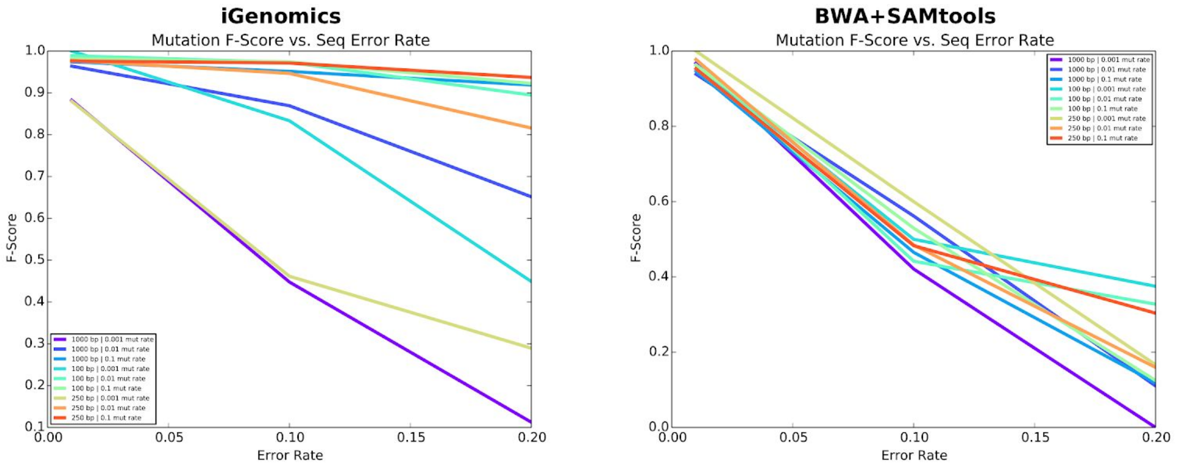


**Supplemental Figure S1.** Mutation identification accuracy for simulated H1N1 flu datasets of varying mutation rates and read length for iGenomics (left) and the BWA-MEM/Samtools (right) pipeline. The results were computed in the same manner as described in Results section 3 *(Simulated accuracy analysis)*: the simulated reads consisted of H1N1 read sets simulated with an average coverage value of 100 and for all combinations of the following sets of parameters: sequence error rates of 0.01, 0.1, and 0.2, mutation rates of 0.001, 0.01, and 0.1, and read lengths of 100bp, 250bp, and 1,000bp.

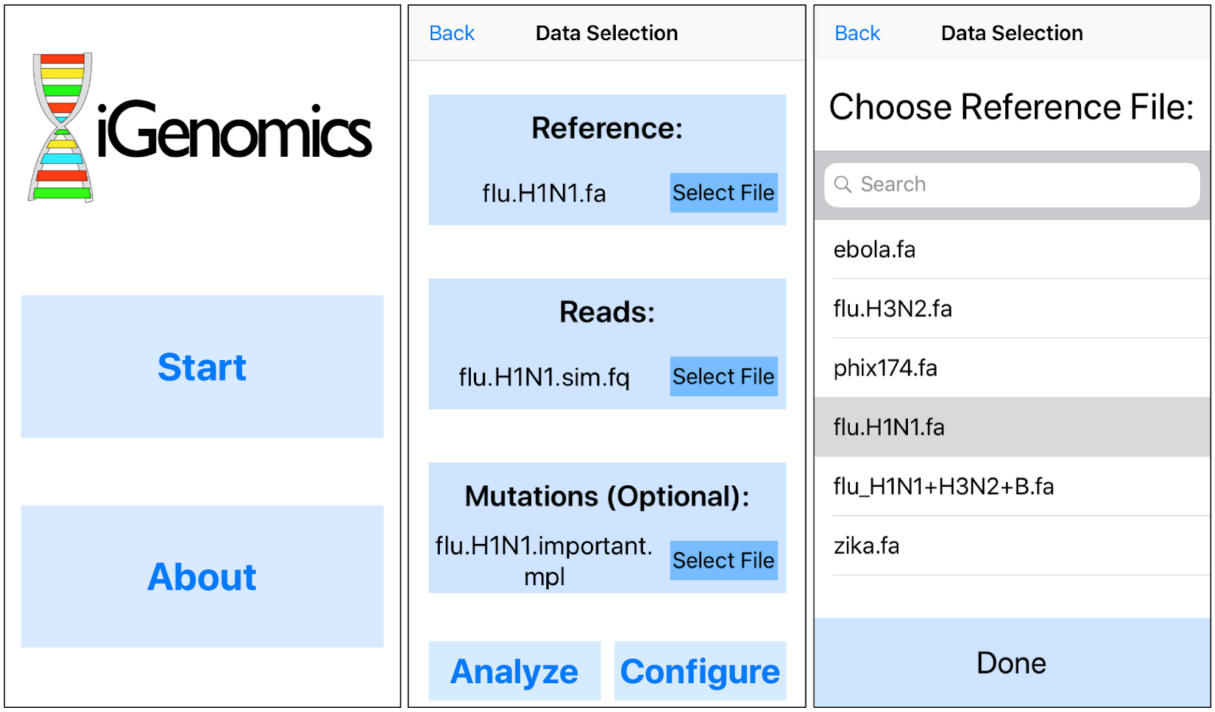


**Supplemental Figure S2.** iGenomics iPhone screenshots: (left) launch screen, (middle) file selection page, (right) individual file selector. By pressing the 'Start' button on the launch screen, the user is brought to the file selection page. Pressing 'Select File' on the file selection page will allow the user to use the individual file selector to choose a default file (pre-packaged with iGenomics) or local file (saved to iGenomics from an external app) or to use Dropbox's UI to choose a file from the user's Dropbox account. Additionally, the user can select 'Analyze', which will immediately begin to align the input reads to the reference using the most recently used parameters, or 'Configure', which will present the parameter selection page.

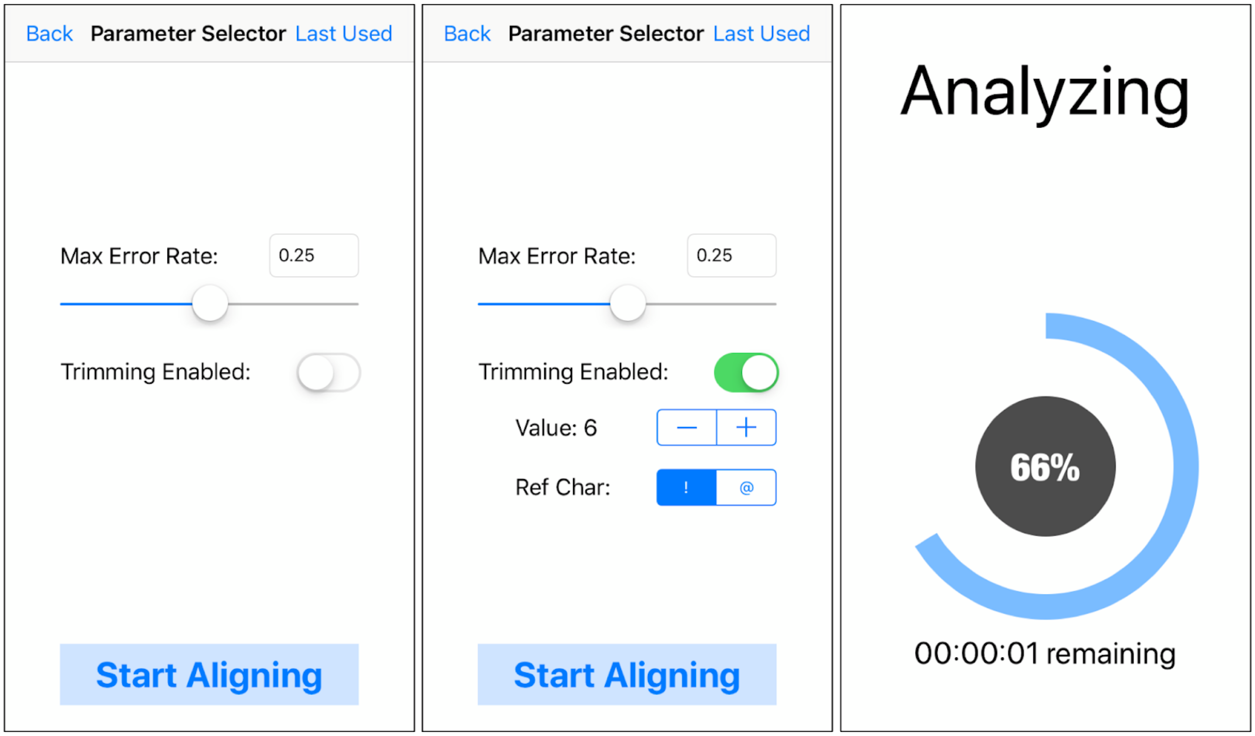


**Supplemental Figure S3.** iGenomics iPhone screenshots: (left) parameter selection page with trimming disabled, (middle) parameter selection page with trimming enabled, (right) computing page. From the file selection page in Supplemental Figure S2, if the user chooses ‘Analyze’, the right computing page will be shown and if the user chooses ‘Configure’, the parameter selection page will be shown with the last used parameters. Pressing ‘Start Aligning’ from the parameter selection page will begin aligning the reads using the configured parameters. On the computing page, the percentage indicates the total percent of reads aligned and the time remaining indicates the estimated time remaining before the alignment and variant identification process completes.

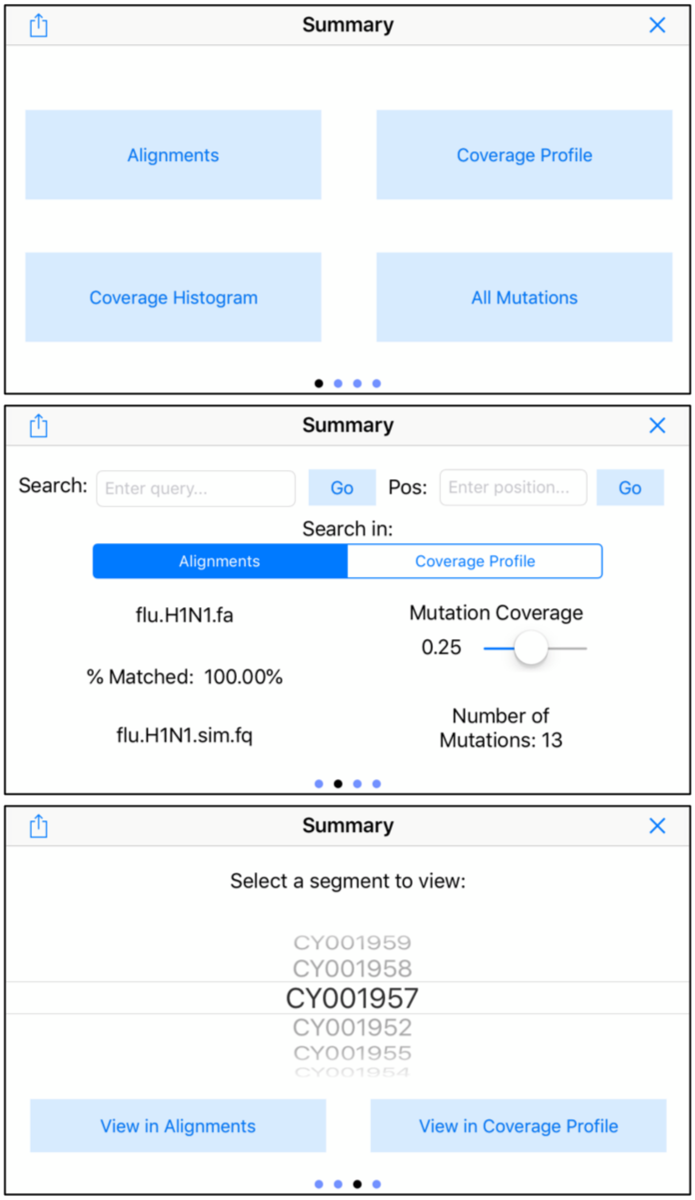


**Supplemental Figure S4.** iGenomics iPhone screenshots: (top) view selection page, (middle) alignment details page, (bottom) segment selection page. The view selection page allows the user to view the alignments display and coverage profile (shown in Figure 2) as well as the coverage histogram and found mutations list (shown in Supplemental Figure S5). The alignment details page displays information about the alignments, including the reads and reference file names, percent of reads that matched, and the number of mutations, and allows the user to search the reference genome and adjust the minimum relative heterozygosity value (known as mutation coverage within iGenomics). The segment selection page lets the user intuitively choose a particular segment in the reference genome to view alignment information for. These three pages, in addition to a fourth page (the important mutations display shown in Figure 1), can be navigated with just a swipe.

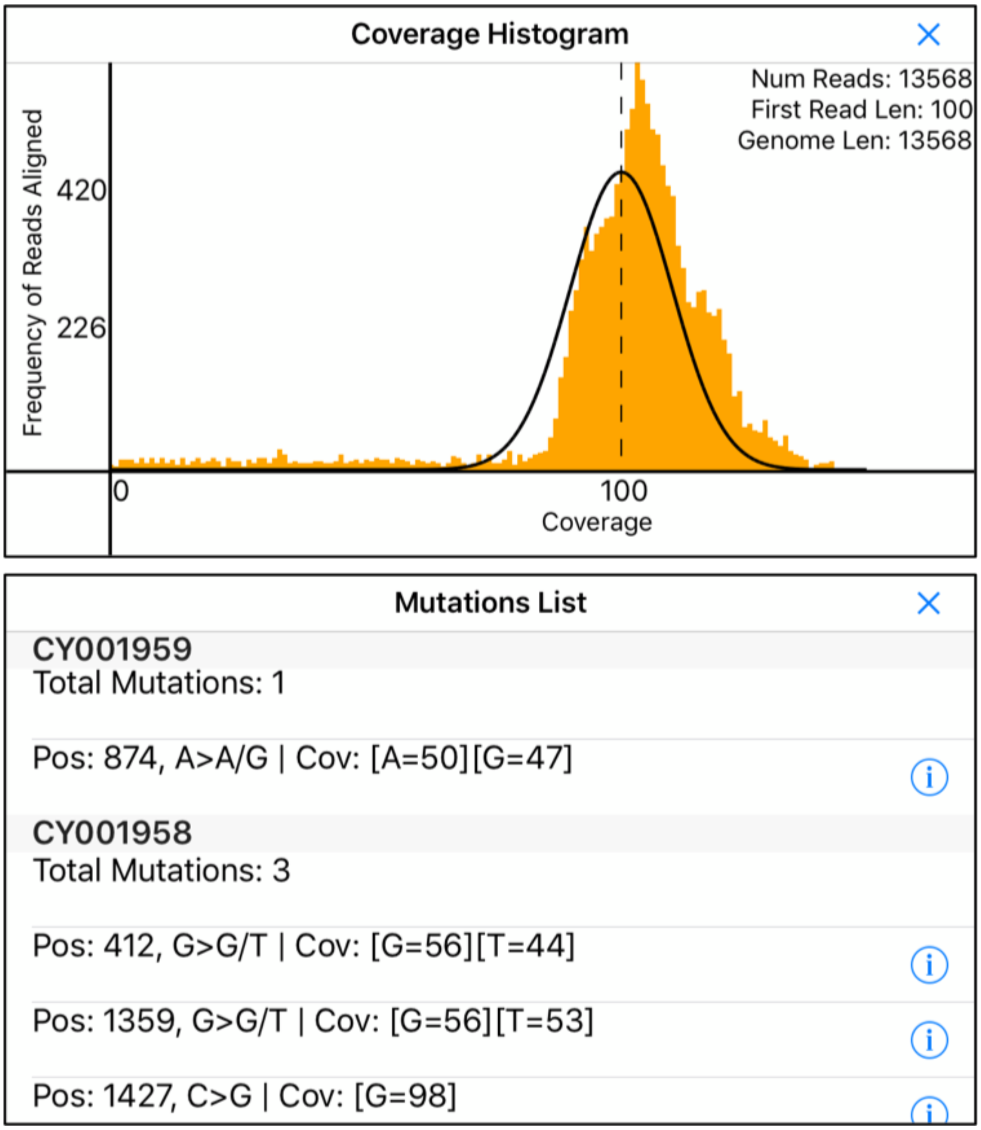


**Supplemental Figure S5.** iGenomics iPhone screenshots: (top) coverage histogram, (bottom) found mutations list. The coverage histogram displays a plot of the frequencies of each coverage value with a poisson curve for context. In these screenshots, we used simulated H1N1 reads of length 100bp with an average coverage value of 100. The found mutations list displays the number of mutations identified in each segment and information about each of those mutations. By tapping the circle `i` icon, the user can navigate directly to the mutation in the coverage profile or alignments view (whichever was most recently used). Adjusting the mutation coverage slider in Supplemental Figure S4 will affect the mutations that are displayed in this list.

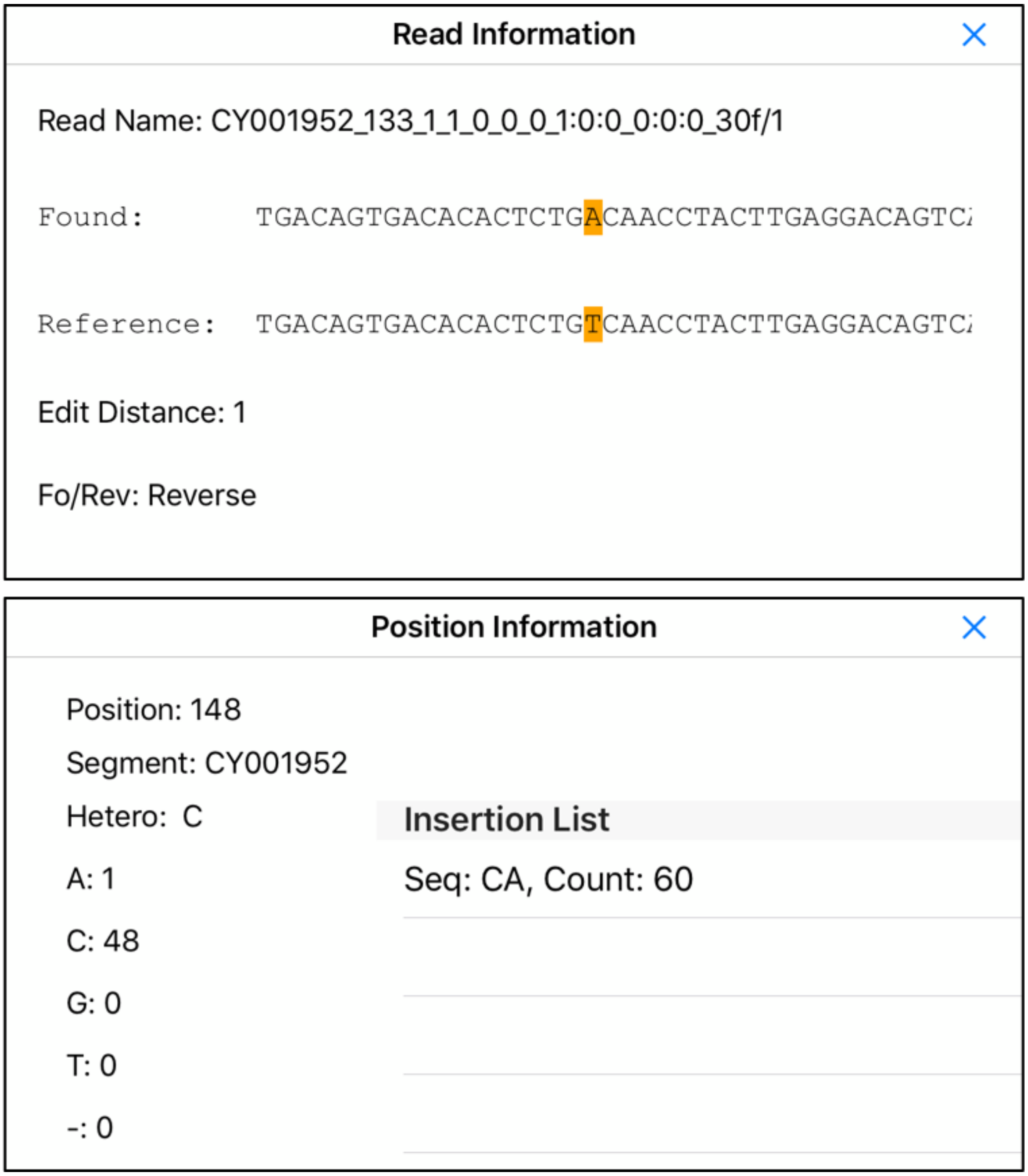


**Supplemental Figure S6.** iGenomics iPhone screenshots: (top) position information popover, (bottom) read alignment popover. The position information popover for a given position displays coverage details, heterozygosity, and, if present, insertion mutations. This popover can be invoked by double-tapping anywhere in the column for a position from within the alignments display or coverage profile. The read alignment popover shows specifically how a particular read aligned to the reference genome, and can be brought up from the alignments display by long pressing an aligned read.

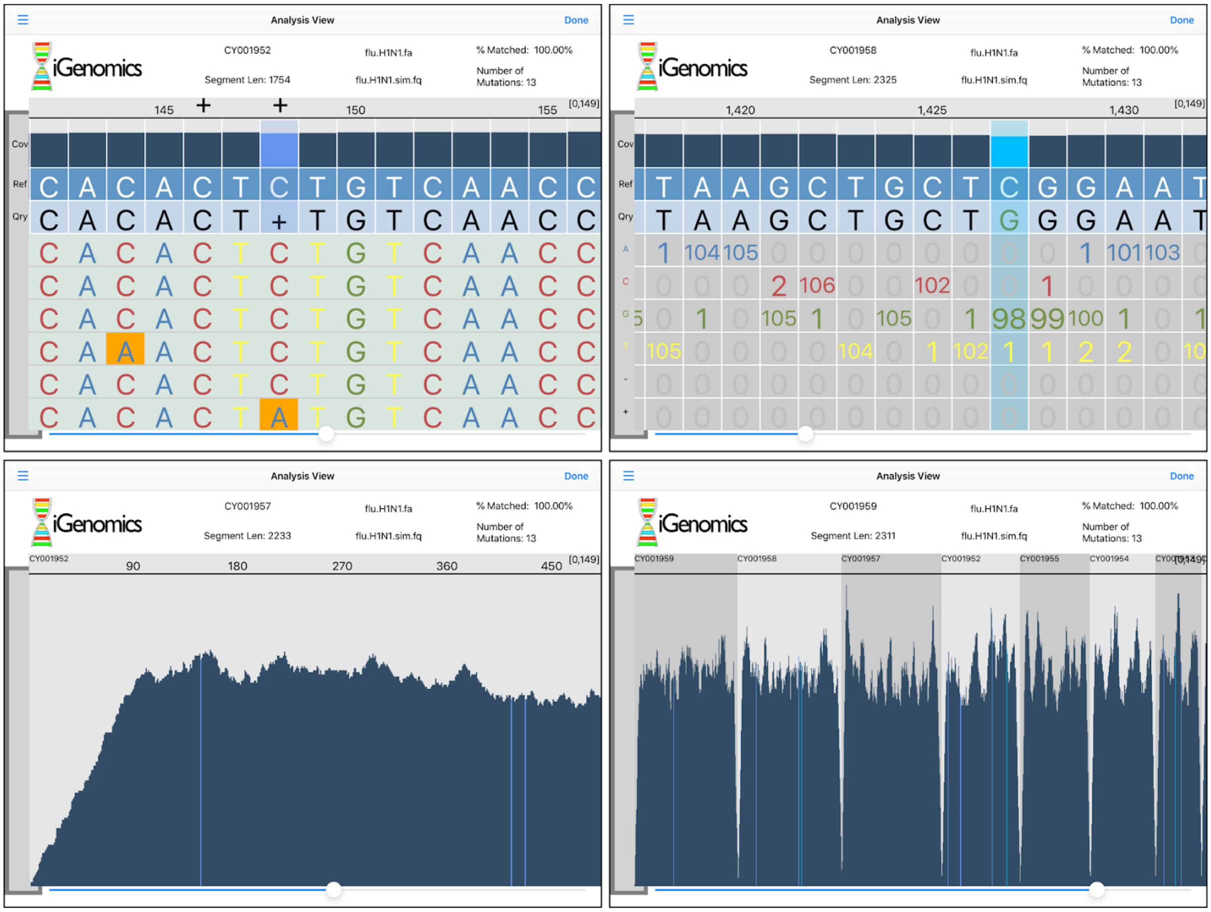


**Supplemental Figure S7.** iGenomics iPad screenshots: (top-left) alignments display, (top-right) coverage profile, (bottom-left) partially zoomed-out coverage profile, (bottom-right) fully zoomed-out coverage profile. The iPad application for iGenomics strongly resembles that of the iPhone application for all views except the analysis ones. In the analysis view, alignment details are always visible at the top of pane and the alignments display/coverage profile is displayed below the details. As with the iPhone version of iGenomics, the user can switch between the alignments display and coverage profile and can zoom out of either to see the relative coverage at varying levels of granularity.

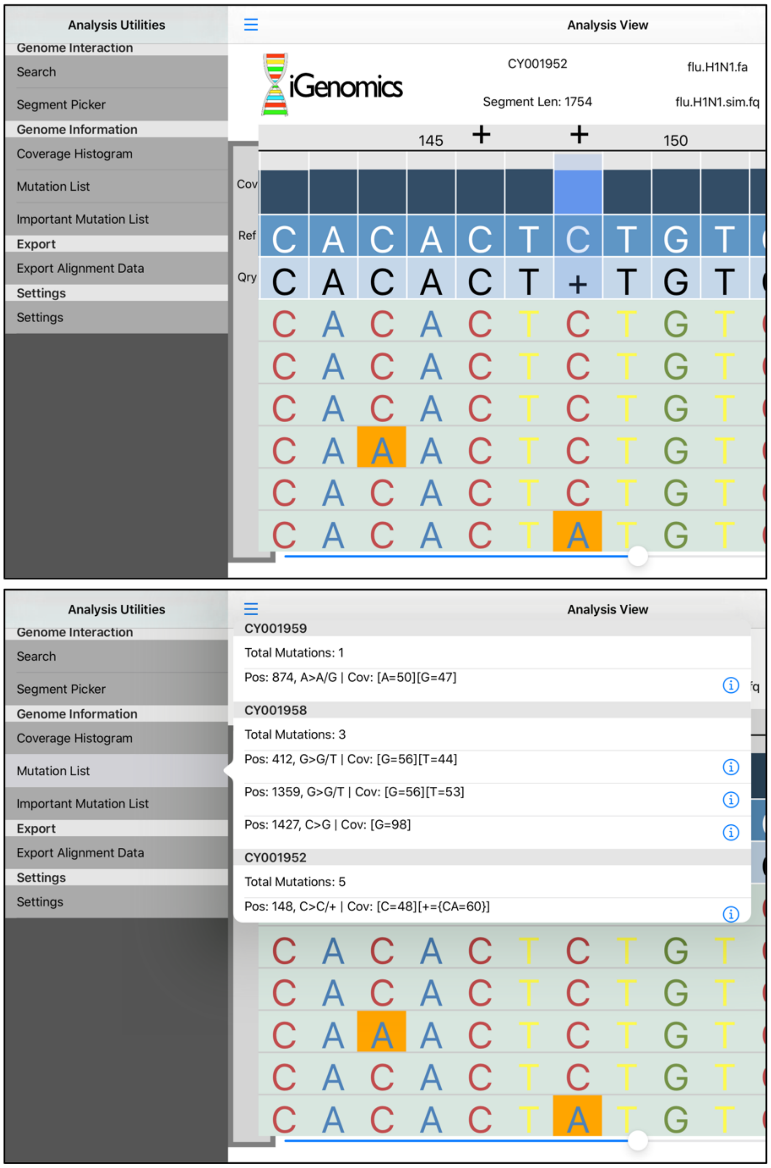


**Supplemental Figure S8.** iGenomics iPad screenshots: (top) analysis utilities, (bottom) found mutations list. Tapping three line icon in the top left of the analysis view will bring up the analysis utilities, which contains the same capabilities as the iPhone version of iGenomics but presents views in iPad-native popovers rather than new fullscreen pages. Tapping on any of these utilities, such as the “Mutation List”, will present the results in a popover.
